# New land use change scenarios for Brazil: refining global SSPs with a regional spatially-explicit allocation model

**DOI:** 10.1101/2021.08.12.456156

**Authors:** Francisco Gilney Silva Bezerra, Celso Von Randow, Talita Oliveira Assis, Karine Rocha Aguiar Bezerra, Graciela Tejada, Aline Anderson Castro, Diego Melo de Paula Gomes, Rodrigo Avancini, Ana Paula Aguiar

## Abstract

The future of land use and cover change in Brazil, in particular due to deforestation and forest restoration processes, is critical for the future of global climate and biodiversity, given the richness of its five biomes. These changes in Brazil depend on the interlink between global factors, due to its role as one of the main exporters of commodities in the world, and the national to local institutional, socioeconomic and biophysical contexts. Aiming to develop scenarios that consider the balance between global and local factors, a new set of land use change scenarios for Brazil were developed, aligned with the global structure Shared Socio-Economic Pathways (SSPs) and Representative Concentration Pathway (RCPs) developed by the global change research community. The narratives of the new scenarios align with SSP1/RCP 1.9, SSP2/RCP 4.5, and SSP3/RCP 7.0. The scenarios were developed combining the LuccME spatially explicit land change allocation modeling framework and the INLAND surface model to incorporate the climatic variables in water deficit. Based on detailed biophysical, socio-economic and institutional factors for each biome in Brazil, we have created spatially-explicit scenarios until 2050, considering the following classes: forest vegetation, grassland vegetation, planted pasture, agriculture, mosaic of small land uses, and forestry. The results aim at regionally detailing global models and could be used both regionally to support decision-making, but also to enrich global analysis.

## 1 Introduction

Land available for agricultural expansion is an increasingly scarce resource in several global regions [1, 2]. The agricultural frontier’s expansion, currently concentrated in the tropics [3, 4], affects the regulation of the hydrological and climatic regime, local socioeconomic relations and generates a great biodiversity loss. This context could be aggravated if we consider population increase and demand for food 2050 projections (25% and 40%, respectively) [5].

In this context, global models and scenarios, particularly the ones quantified with Integrated Assessment Models (IAMs), that represent complex interactions and feedback on a long term scale between the socioeconomic system (including climate policies) and the natural system [6], playing a key role in helping us to understand the impacts and consequences of agricultural expansion in different regions. In Brazil, for example, this process over the last few decades has contributed to the country consolidating worldwide as one of the main commodity-exporting countries, whether agricultural or mineral. One of the key impacts of this process is the loss of natural vegetation in the Amazon and Cerrado biomes. On the other hand, other areas in Brazil, such as the Atlantic Forests, are undergoing a forest transition process [7]. The integration and understanding of the factors that influence land use and land cover changes (LUCC) in Brazil in different regions are important in order to define indicators for guiding public policies to establish sustainable development strategies.

Global models and scenarios may fail to capture the regional land change dynamics as they do not always include local factors, regional narratives, the national political and institutional structure, and the dynamics and magnitude of intraregional drivers, which determine the demand for land. In addition, the information used in most global models is aggregated for comparability between large regions, such as continents, etc. [8–10]. In this sense, Dala-Nora [8] points out that a balance between global and local factors is necessary since the integration between these complex factors that operate on a global and regional scale through extensive flow networks can change the structure and consistency of land use change scenarios. In addition,VanVuuren [11] points out that studies that examine phenomena of a more precise scale should consider more detailed information (eg, geographic characteristics, land use patterns, or the location of cities).

In this paper, we present a new set of land change scenarios for Brazil, aligned with the global Shared Socio-economic Pathways (SSPs) and Representative Concentration Pathway (RCPs) framework developed by the global change research community [9, 12–16] to support the Intergovernmental Panel on Climate Change (IPCC). These scenarios are being widely used and downscaled to several regions (eg, [16–20]) and adopting them as a reference allows us to better link our scenarios to the global context. The regionalized scenarios developed here aim to represent the diversity of processes linked to land use change in the Brazilian territory. The modeling approach considers Brazilian biomes’ interregional socio-ecological differences, including a more detailed analysis scale, without losing their relationship with global relations. We model changes in natural vegetation, large and small-scale agricultural lands, and planted forests. These land change processes are directly related to regional and local factors, and the global context still plays a significant role in these processes.

## 2 Materials and methods

### 2.1 Overall conceptualization and structure

The **regional scenarios** were quantified by the LuccME modeling framework [21]. LuccME is a spatially explicit dynamic modeling structure for LUCC developed at the Institute for Space Research - INPE. This approach makes it possible to delineate the spatial patterns of land use and land cover classes based on the components of (a) **Demand**, that is the amount/intensity of changes in each use that is intended to be allocated over time [22, 23]; (b) **Potential**, which corresponds to the adequacy that a given cell in the cellular space has to change with each step of time, complete, in this case, using the Spatial Lag regression model [23–25] and (c) **Allocation** that spatially and interactively distributes the LUCC according to the previous components (demand and potential), based on competition between classes of land use in each cell. The land use and cover data used in the LuccMEBR comes from IBGE [26]. We chose this database due to its national scope, periodicity (2000, 2010, 2012, and 2014), and classes. Also, IBGE data is consistent with other regional mappings as TerraClass http://www.inpe.br/cra/projetos_pesquisas/dados_terraclass.php and MapBiomas https://plataforma.brasil.mapbiomas.org/.

For **global scenarios**, we use the projections generated by the Integrated Model to Assess the Global Environment (IMAGE) [27, 28], representing different combinations of SSPs and RCPs. The integration across scales to generate the scenarios happens as follows. First, we defined the land change classes. Table 1 presents the description of the original and reclassified IBGE classes used for the LUCC modeling and the equivalent classes in the IMAGE model. The thirteen classes from IBGE [26] were reclassified, grouping the similar classes to reduce the complexity of the model. Second, we used the quantity of change projected by IMAGE for the selected combinations of SSPs/RCPs to define the quantity of change for each land use (LuccME demand component), as detailed in Section 2.3. Third, we developed local narratives allied to the selected SSPs/RCPs. Based on these narratives, we parameterized the LuccME allocation and potential components using a comprehensive database of socioeconomic, institutional, and biophysical drivers. Fourth, climate projection data for some of the RCPs were used to generate water deficit data using the Integrated Surface Process Model (INLAND) model. Fig 1 illustrates the integration/translation structure across scales to generate regionalized scenarios.

**Table 1.** Land use and land cover classes.

| Original land use and cover classes - IBGE | LuccME/BR reclassification | Original land use and cover classes - IMAGE | Description |
| --- | --- | --- | --- |
| Forest vegetation<br>Humid Areas | Forest vegetation | Tropical woodland<br>Tropical forest | Areas occupied by forests, which include areas of Dense Forest (forest structure with continuous top cover), Open Forest (forest structure with different degrees of discontinuity of the top cover, according to its type with vine, bamboo, palm or sororoca), Forest Seasonal (forest structure with loss of leaves from the upper strata during the unfavorable season - dry and cold), in addition to the Mixed Rain Forest (forest structure that comprises the natural distribution area of Araucaria angustifolia, a striking element in the upper strata, which generally forms a cover to be continued). In addition to these, other features were included, such as Savannah Forest, Campinarana Forest, Campinarana Arborizada, and Mangroves, as well as humid areas, which correspond to natural herbaceous vegetation, permanently or periodically flooded with fresh or brackish water (estuaries, swamps, etc.). In these areas the land of ponds, swamps, humid fields, are inserted, among others. |
| Natural pasture | Grassland vegetation | Extensive grassland<br>Scrubland<br>Savana | The area is occupied by grassland vegetation subject to grazing and other low-intensity anthropic interference. |

Table 1 – *Table continuation*
| Original land use and cover classes - IBGE | LuccME/BR reclassification | Original land use and cover classes - IMAGE | Description |
| --- | --- | --- | --- |
| Pasture planted | Pasture planted | Grassland-steppe | The area is predominantly occupied by cultivated herbaceous vegetation. They are places for cattle grazing, formed by planting perennial forages, subject to high-intensity anthropic interference, such as clearing the land (unblocking and cutting), liming, and fertilizing. |
| Agriculture | Agriculture | Agricultural land<br>Biofuels | Areas occupied by temporary crops and permanent crops, irrigated or not, being land used for the production of food, fiber and agribusiness commodities. It includes all cultivated land, which may be planted or at rest, and also as a cultivated wetland. It can be represented by heterogeneous agricultural zones or extensive areas of plantations. |
| Mosaic of agricultural area with forest remnants | Mosaic of occupation | Warm mixed forest | Mosaic of small-scale agricultural activities and remnants of natural vegetation (eg, subsistence agriculture). |
| Mosaic of Forest Vegetation with Agricultural Activity |  |  |  |
| Mosaic of agricultural area with grassland remnants |  |  |  |
| Forestry | Forestry | Regrowth forest timber | Forests planted and managed with exotic species (eg eucalyptus, pine). |
| Artificial Area | Others | Not considered | Artificial surfaces (eg cities, villages, transport routes, energy and communication networks) and bodies of water and bare lands (eg rock outcrops, dunes, cliffs). |
| Bare land |  |  |  |
| Inland water bodies |  |  |  |
| Coastal water bodies |  |  |  |

**Fig 1.**
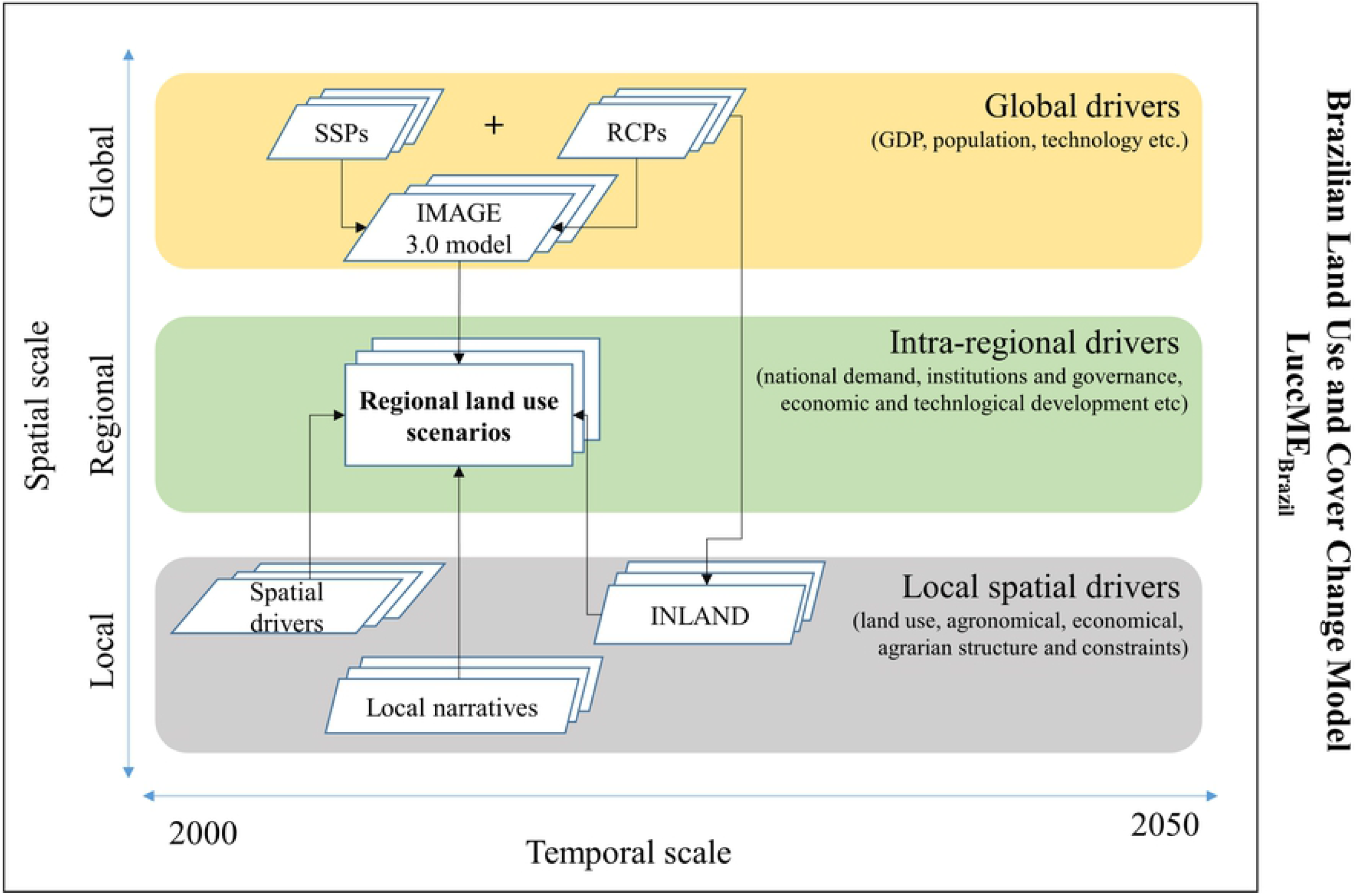
Schematic representation of the development of regional land use and land cover scenarios. Shared Socio-Economic Pathways (SSPs) and Representative Concentration Pathway (RCPs).

### 2.2 Scenario assumptions: from global to regional

The SSPs are based on five different development paths for societal trends (Table 2: i.e., sustainable development (SSP1), middle of the road developments (SSP2), global fragmentation (SSP3), strong inequality (SSP4), and rapid economic growth based on a fossil-fuel intensive energy system (SSP5). They were designed to represent different degrees of challenges to mitigation and adaptation. Each of the SSPs has been elaborated in a storyline and quantified using Integrated Assessment Models (IAM), such as IMAGE. The five SSP storylines can also be combined with alternative assumptions about climate mitigation, forming a matrix of alternative scenarios.

**Table 2.** Synthesis of core premises differentiating the SSPs in relation to land use. Source: (Popp et al., 2017).

|  | SSP 1 | SSP 2 | SSP 3 | SSP 4 | SSP 5 |
| --- | --- | --- | --- | --- | --- |
| Land-use change regulation | Strong regulation to avoid environmental tradeoffs | Medium regulation; slow decline in the rate of deforestation | Limited regulation; continued deforestation | Highly regulated in medium-income (MICs) and high-income (HICs) countries; lack of regulation in low-income countries (LICs) lead to high deforestation rates | Medium regulation; slow decline in the rate of deforestation |
| Land productivity growth | High improvements in agricultural productivity; rapid diffusion of best practices | Medium pace of technological change | Low technology development | High productivity for large scale industrial farming, low for small-scale farming | Highly managed, resource-intensive; rapid increase in productivity |
| Environmental Impact of food consumption | Low growth in food consumption, low-meat diets | Material-intensive consumption, medium meat consumption | Resource-intensive consumption | Elites: high consumption lifestyles; Rest: low consumption | Material-intensive consumption, meat-rich diets |
| International Trade | Moderate | Moderate | Strongly constrained | Moderate | High, with regional specialization in production |
| Globalization | Connected markets, regional production | Semi-open globalized economy | De-globalizing, regional security | Globally connected elites | Strongly globalized |
| Land-based mitigation policies | No delay in international cooperation for climate change mitigation. Full participation of the land use sector | Delayed international cooperation for climate change mitigation. Partial participation of the land use sector | Heavily delayed international cooperation for climate change mitigation. Limited participation of the land use sector | No delay in international cooperation for climate change mitigation. Partial participation of the land use sector | Delayed international cooperation for climate change mitigation. Full participation of the land use sector. |

Therefore, each SSP has a baseline scenario implementation and additional scenarios combining the storyline to climate mitigation policies compatible with certain levels of CO2 concentration - and consequently, climate change. These mitigation assumptions are linked to RCPs, meaning the atmosphere’s expected radiative forcing in 2100 (W/m2). For instance, the RCP 2.6 assumes a 3 W/m2 peak (490 ppm CO2 eq) before 2100 and a decline to 2.6 W/m2 by 2100.

We have adopted in our regionalized scenarios the following combinations: SSP1 RCP 1.9, SSP2 RCP 4.5 and SSP3 RCP 7.0 (Table 3). In particular, the sustainable development scenario (SSP1) combined with stringent climate policy (RCP 1.9) is a scenario exploring the route towards a more sustainable world - providing an initial framework for our analysis of sustainability pathways. Although the Sustainable Development Goals (SDGs) were not targeted in its development ([20]). Mitigation scenarios that achieve the ambitious targets included in the Paris Agreement typically rely on greenhouse gas emission reductions combined with net carbon dioxide removal from the atmosphere, mostly accomplished through the large-scale application of bioenergy with carbon capture and storage, and afforestation (eg, [16, 28]).

**Table 3.**
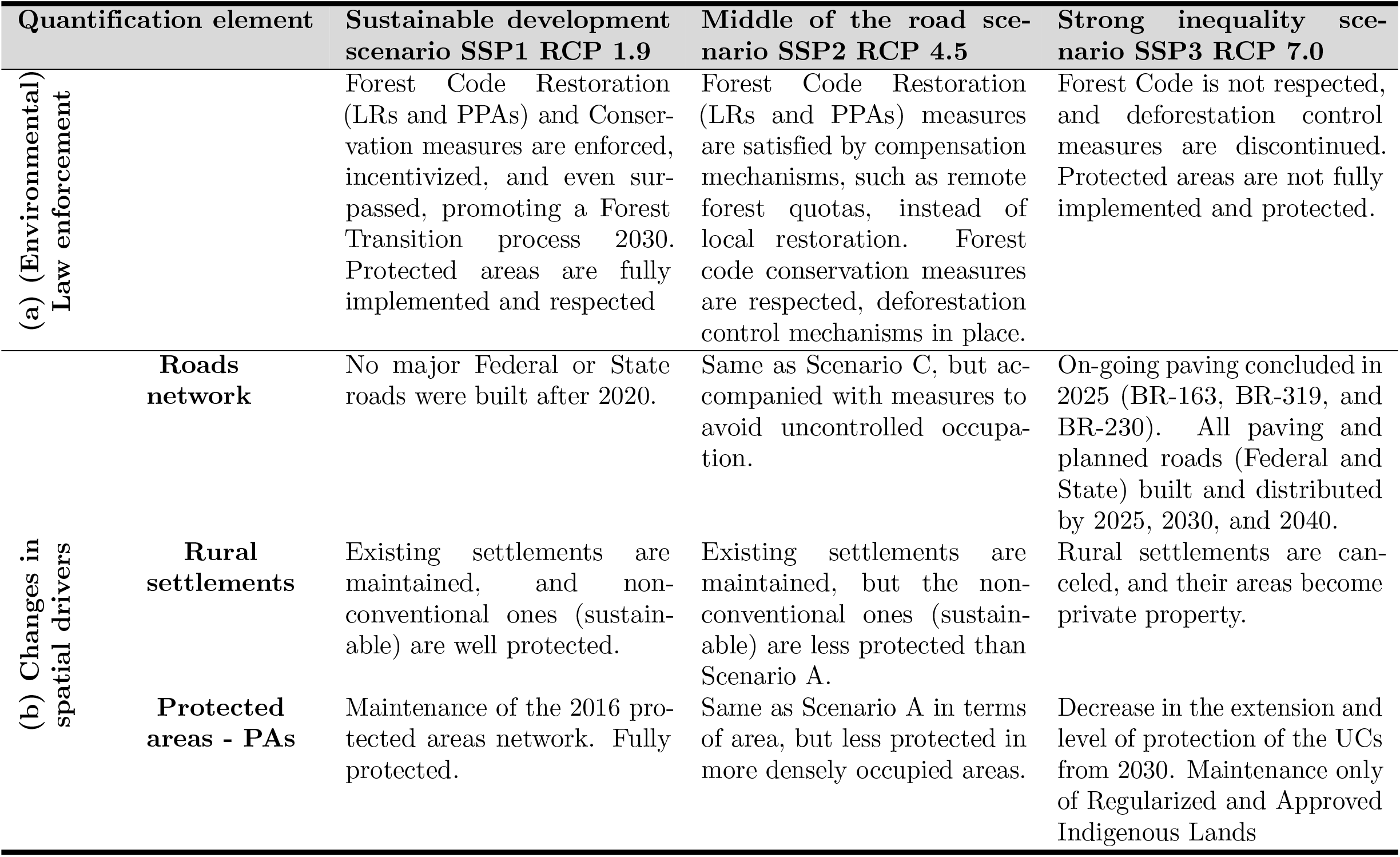
Detailed regional assumptions related to Sustainable development scenario, Middle of the road, and Strong inequality scenario.

### 2.3 LuccME model parameterization and validation

#### 2.3.1 Demand component

We use the LuccME ”PreComputedValues” component, in which we externally calculate demand and report to the model the expected area for each land use class annually, in the period 2000 to 2050 (Tabela 4). As described above, we use the amount of change projected by IMAGE and adjusted to the IBGE land use and land cover classes for the SSP1/RCP1.9, SSP2/RCP4.5, and SSP3/RCP7.0 combinations, to generate the annual demand for each land use class in each scenario, between 2015 to 2050.

Equation 1 presents the calculation of the annual change ***C***_***ca***_ for each class of land use and land cover in area unit:

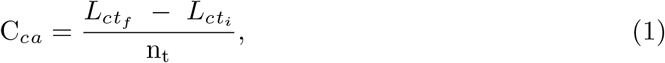

where ***C***_***ca***_ corresponds to the annual change, in area, of the land use class ***L***_***c***_ between initial ***t***_***i***_ and ***t***_***f***_ end year of the chosen period; and ***n***_***t***_ refers to the number of years in the period.

The calculation of the annual demand 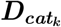 of the land use class is represented by equation 2:

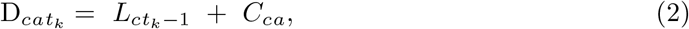

where 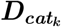 corresponds to the annual demand, in area, of a given land use class ***L***_***c***_ in a given year ***t***_***k***_, calculated from the sum of the class area in the previous year ***t***_***k***−***1***_ and the annual change ***C***_***ca***_.

In the initial year, the value of the demand corresponds to the observed amount of the land use class, calculated based on the use and land cover data used, in this case, the land use and land cover change data from IBGE [**?**].

#### 2.3.2 Potential and allocation component parametrization: Intra-regional and local spatial drivers

The LuccME component is used to determine the potential occurrence of a given land use cover class, as well as the ***PotentialCSpatialLagRegression*** (Equation 3), which is based on and adapted from the spatial regression model (Spatial Lag) [23–25]). In this component, it is considered that the influence of the neighboring areas occurs. This is an intrinsic feature of land use and land cover change. In addition, this component allows this potential to be dynamic over the modeled period, that is, every year.

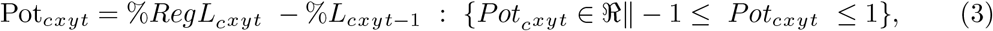

where ***Pot***_***cxyt***_ corresponds to the potential for the occurrence of a given land use class ***L***_***c***_ in a given location ***xy*** in a given time step ***t***. To determine the potential, the percentage of use estimated by the regression ***RegL***_***cxyt***_ is subtracted from the percentage of existing use ***L***_***cxy***_ at time ***t-1***.

For the calculation of the potential, the variables potentially explaining the process of changing land use and coverage in different Brazilian biomes were divided into four categories: agronomic aspects (composed of climatological and geophysical variables); agrarian structure (percentage of the area of agricultural establishments); economic aspects (dependent on structural variables: distance from highways, ports, airports, railways, urban centers, rivers, etc.), and restrictive aspects (related to legal limitations: protected areas, conservation units, rural settlements, distance to hydroelectric and thermoelectric in operation). All candidate variables were integrated into regular 10km x 10km cells for the spatially explicit analysis, taking as base years 2000 and 2010 (Fig. 2). For that, the ***FillCell plugin*** was used [29]. The use of cellular space made it possible to homogenize the factors described above, regardless of their origin format (vector data, matrix data, etc.), aggregating them in the same space-time basis, through operators (for example, Percentage of each class, Minimum distance, etc.) used according to the geometric representation and the semantics of the attributes of the input data.

**Fig 2.**
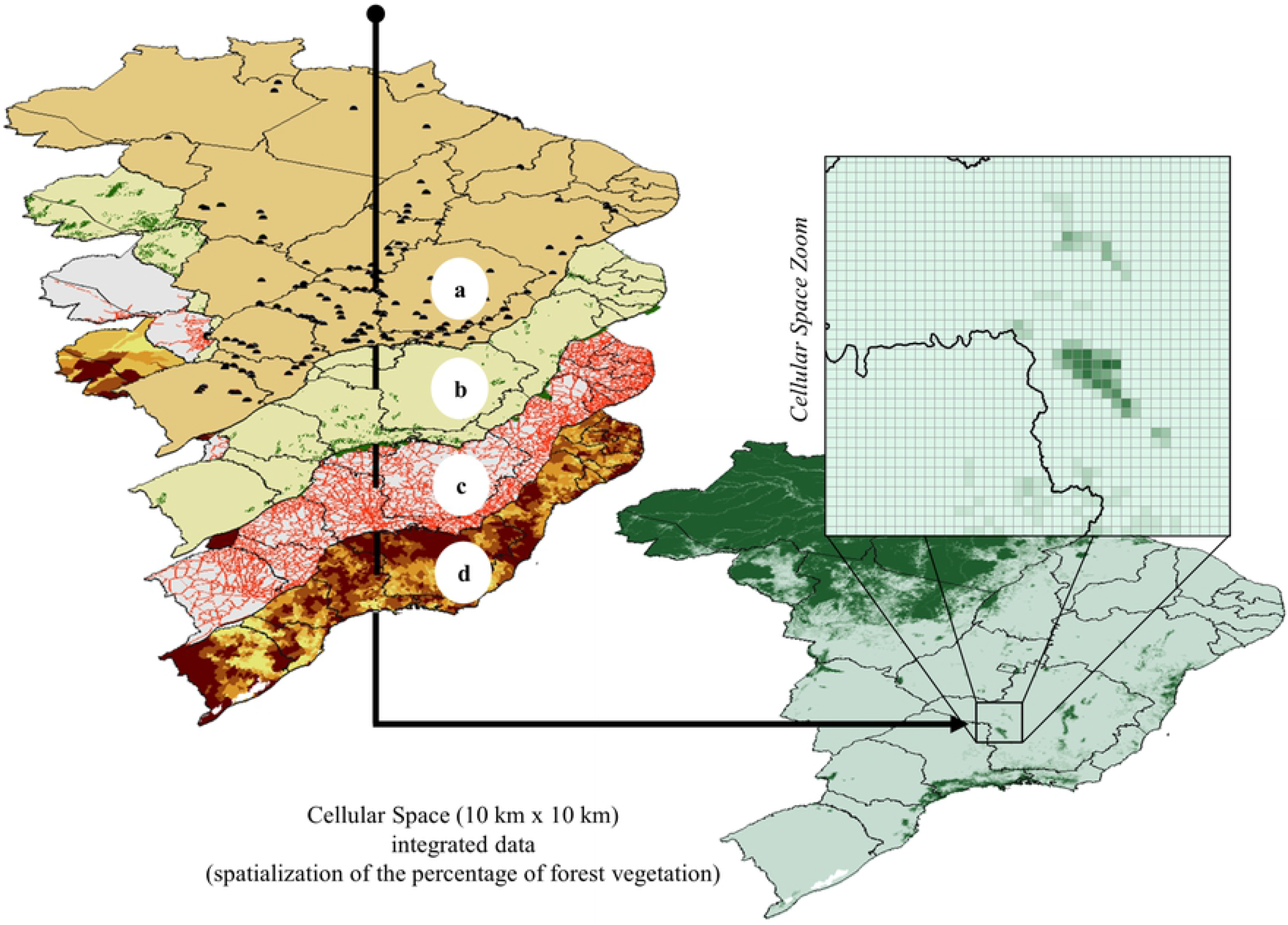
Representation of the factors integration into the cellular space. a) Hydroelectric plants, b) Protected areas, c) Federal and state highways, and d) Proportion of large agricultural establishments.

Searching for a model that involves the minimum of possible parameters to be estimated and explains well the behavior of land use in each Brazilian biome, some statistical techniques were used to evaluate the effectiveness and adequacy of the best model: a priori, the Spearman’s correlation analysis selected only those factors that presented a correlation coefficient below or equal to 0.60; secondly, with the new composition of candidate variables, a spatial regression analysis was performed (Spatial Lag [24, 25]), considering the determination coefficient (*R*^2^ > 0.75, in the average of uses) as decision parameters; statistical significance (p-value < 0.05) and Akaike information criterion - AIC (lowest values obtained). The calibration of the model was carried out using observational data contained in the land use and land cover maps of IBGE in the years 2000, 2010, 2012, and 2014.

The allocation of land use change in each scenario was based on the application of the LuccME AllocationCClueLike allocation component based on the CLUE [30] which was distributed spatially and interactively according to the previous components (demand and potential), based on competition between the types of land use in each cell and within a previously established maximum error. For the parameterization of this component, the Forest Code rules regarding the amount of legal reserve required according to the regions of the Brazilian territory were also considered. The allocation process for each type of land use or land cover can be described using equation 4.

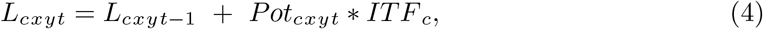

where the amount of area allocated from a given class of land use ***L***_***c***_ at a given ***xy*** location in the cell plane at time ***t*** is determined in an iterative process of the sum of ***L***_***cxy***_ at time ***t*-*1*** and the potential ***Pot***_***cxyt***_, multiplied by an adjustment factor proportional to the difference between the allocated area, the reported demand, and the direction of the change ***ITF***_***c***_.

The parameterization details according to Potential (PotentialCSpatialLagRegression), Allocation (AllocationCClueLike) and Demand (DemandPreComputedValues) components are presented in the supplementary material S1 Appendix and S2 Appendix.

#### 2.3.3 Model validation

The results of the simulations were validated by the multi-resolution adjustment validation metric, adapted from Costanza (1989). This metric allows establishing the level of similarity between the simulated and real maps in different resolutions through sampling windows that increase with each step of time. The calculation of the similarity level can be described according to equation 5:

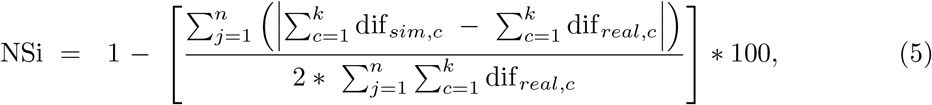

where ***NS*** corresponds to the level of similarity between the real and simulated maps at a given resolution ***i*** ; ***j*** is the window considered; ***n*** establishes the number of windows to be considered; textit**c** is the number of cells in a resolution ***k*** *(i*i)*; and ***dif***_***real***_ = % real_tf_ - % real_ti_ and ***dif***_***sim***_ = % sim_tfinal_ - % real_initial_ being ***ti*** and ***tf*** the initial and real years, respectively, considered in the validation.

## 3 Data Records

The data set provides 35 maps of the spatial distribution of land use and coverage for Brazil between 2000 and 2050, considering the scenarios discussed above. The data are made available in the SIRGAS2000 Polychronic Projection System with Datum (EPSG 5880). The file is available in Zenodo (https://zenodo.org/record/5123560) [31] in NetCDF formats. The data set contains the maps with the percentage of classes Forest vegetation, Grassland vegetation, Managed pasture, Agriculture, Mosaic of occupation, and Forestry in 10 km × 10 km cells.

## 4 Model performance

The performance of the LuccMEBR model was satisfactory, with an average index of spatial adjustment between the observed and simulated of 88% when comparing the patterns of both maps (Fig 3). When considering only the areas where some change process occurred, the adjustment index was 61% for each land use type. The average percentage of adjustment errors corresponding to the omissions was 15%, while the commission errors were approximately 11%.

**Fig 3.**
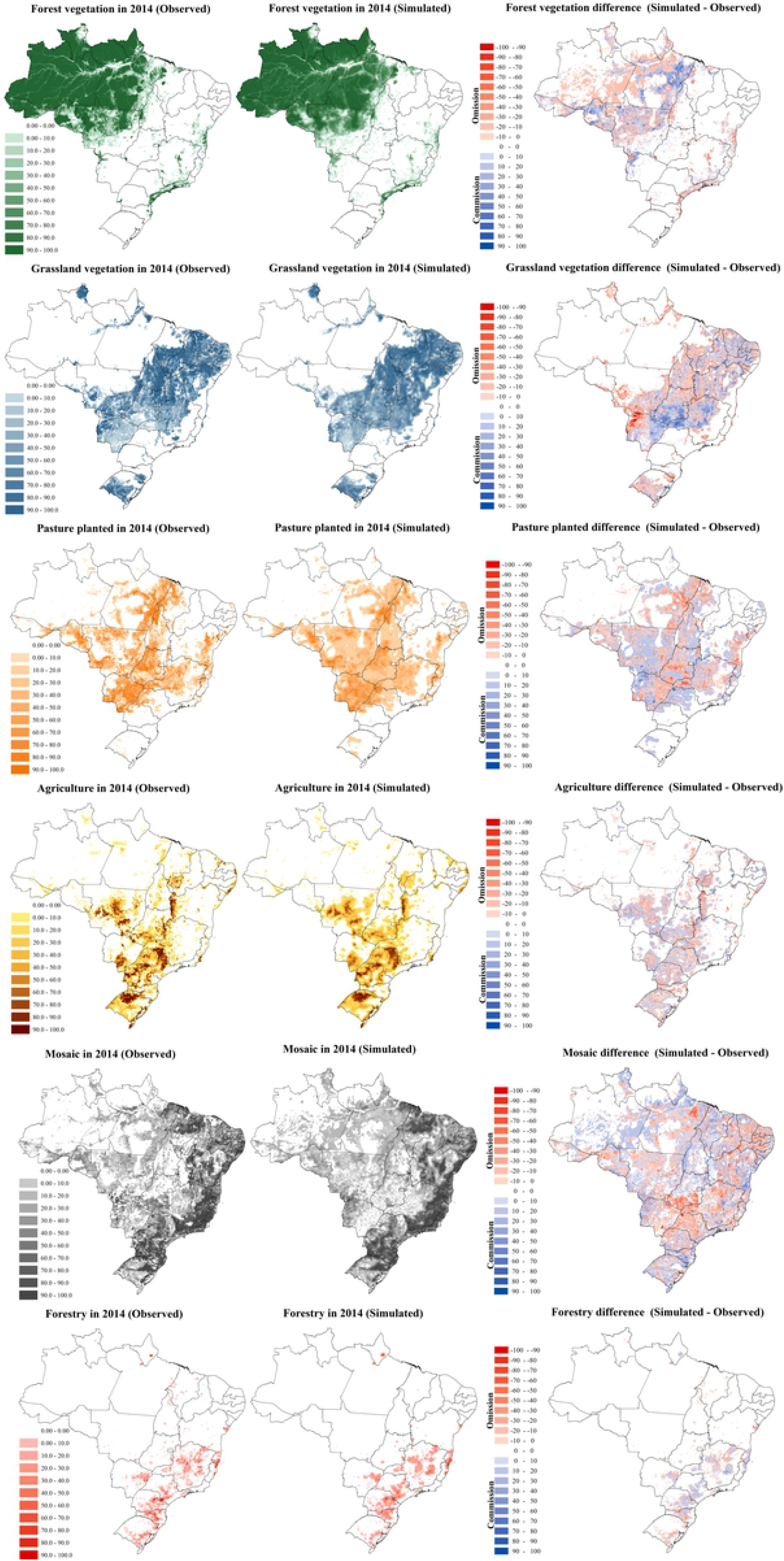
Percentage of each class of land use observed versus simulated in 10 × 10 km^2^ cells in 2014 and the spatial distribution of errors of omission and commission.

## 5 Usage notes

The data on land use and coverage presented for the entire Brazilian territory comes from an effort to align the development of regional scenarios with the structure of global scenarios (RCP-SSP-SPA) developed by the IPCC. Despite the important challenges that this alignment adds to the development of these scenarios, which are: (i) the additional complexity in capturing the multiple dimensions of change and (ii) the issues of scale [32], the result obtained from this process has greater consistency between the different spatial and temporal scales of interest. In addition, the development and study of regional scenarios help policymakers and the scientific community to develop robust strategies in the face of uncertain futures, evaluating and improving the feasibility, flexibility, and concreteness of their actions [33–37]. Fig 4 shows the spatially explicit distribution of the classes of use in the initial year of the simulation (2000) and the three scenarios considered. It can be seen that the Middle of the road and Strong inequality scenarios present similar patterns in all classes of use, with emphasis on the significant increase in the mosaic class of occupations, with intensification in the Caatinga and Atlantic Rainforest. Unlike the other scenarios, the regeneration of forest vegetation is observed in the Sustainable development scenario, mostly in the Caatinga, Atlantic Rainforest, Cerrado, and Pampa biomes.

**Fig 4.**
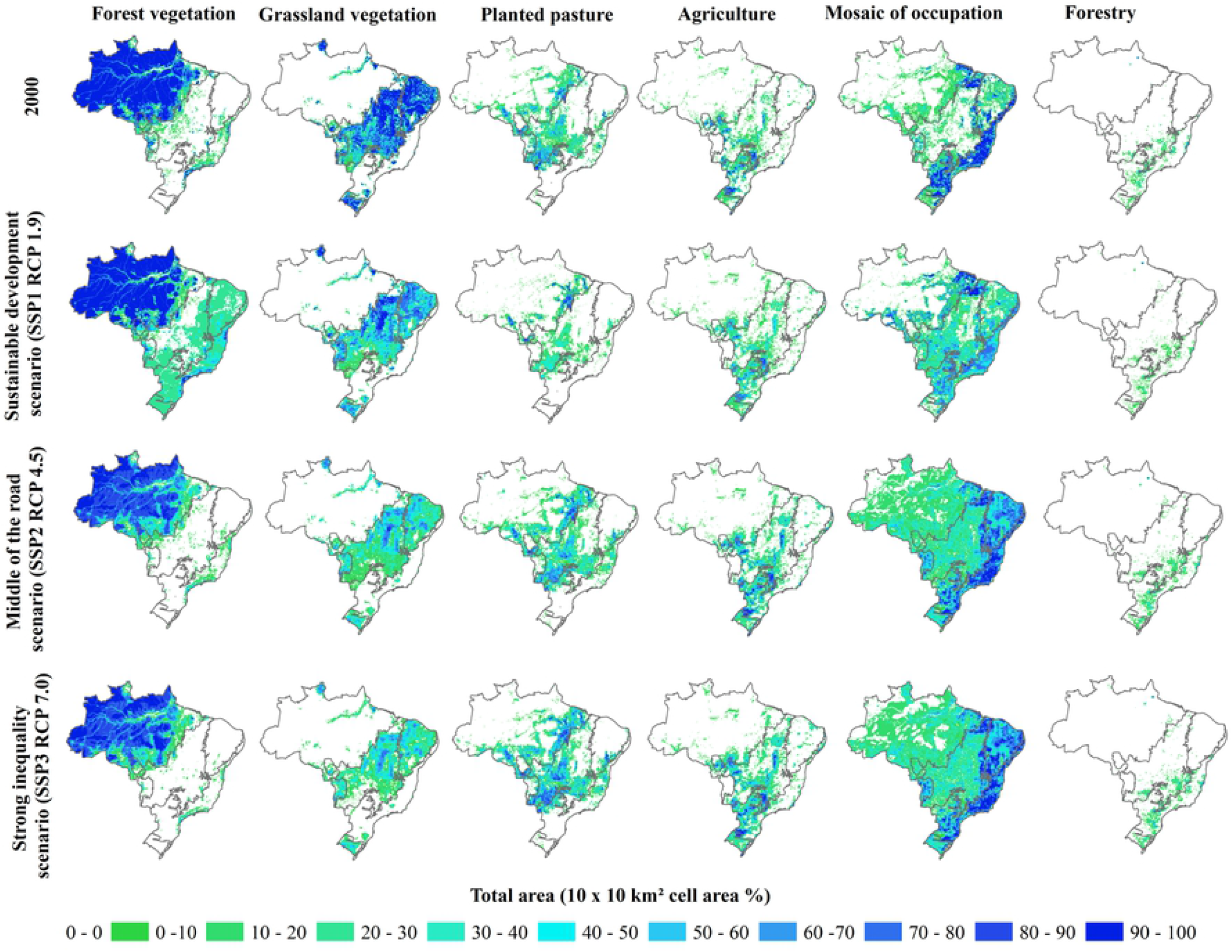
Spatial distribution of areas and land use according to the scenarios from 2000 to 2050.

Analyzing the dynamics of LUCC (Table 5), according to the scenarios considered, has been observed, Agriculture, the Mosaic of occupation, as well as Grassland vegetation, will continue in the same direction, regardless of the scenario considered. In relation to the other classes, it can be seen that the Sustainable development scenario is distinguished from the others, as well as in the spatial pattern observed in Fig 4.

**Table 4.** Land Demand parameters.

| PreComputed values |  | SSP1 RCP 1.9 |  | SSP2 RCP 4.5 |  | SSP3 RCP 7.0 |  |
| --- | --- | --- | --- | --- | --- | --- | --- |
| Forest vegetation | from | 3,771,677 | km<sup>2</sup> (2000) | 3,771,677 | km<sup>2</sup> (2000) | 3,771,677 | km<sup>2</sup> (2000) |
|  | to | 4,232,402 | km<sup>2</sup> (2050) | 2,975,314 | km<sup>2</sup> (2050) | 2,846,765 | km<sup>2</sup> (2050) |
| Grassland vegetation | from | 2,112,947 | km<sup>2</sup> (2000) | 2,112,947 | km<sup>2</sup> (2000) | 2,112,947 | km<sup>2</sup> (2000) |
|  | to | 1,517,633 | km<sup>2</sup> (2050) | 1,012,637 | km<sup>2</sup> (2050) | 772,878 | km<sup>2</sup> (2050) |
| Agricultural area | from | 624,941 | km<sup>2</sup> (2000) | 624,941 | km<sup>2</sup> (2000) | 624,941 | km<sup>2</sup> (2000) |
|  | to | 328,444 | km<sup>2</sup> (2050) | 856,018 | km<sup>2</sup> (2050) | 1,077,091 | km<sup>2</sup> (2050) |
| Pasture management | from | 402,529 | km<sup>2</sup> (2000) | 402,529 | km<sup>2</sup> (2000) | 402,529 | km<sup>2</sup> (2000) |
|  | to | 445,974 | km<sup>2</sup> (2050) | 670,850 | km<sup>2</sup> (2050) | 714,860 | km<sup>2</sup> (2050) |
| Mosaic of occupations | from | 1,405,301 | km<sup>2</sup> (2000) | 1,405,301 | km<sup>2</sup> (2000) | 1,405,301 | km<sup>2</sup> (2000) |
|  | to | 1,822,044 | km<sup>2</sup> (2050) | 2,788,498 | km<sup>2</sup> (2050) | 2,873,629 | km<sup>2</sup> (2050) |
| Forestry | from | 55,983 | km<sup>2</sup> (2000) | 55,983 | km<sup>2</sup> (2000) | 55,983 | km<sup>2</sup> (2000) |
|  | to | 26,881 | km<sup>2</sup> (2050) | 70,060 | km<sup>2</sup> (2050) | 88,154 | km<sup>2</sup> (2050) |

**Table 5.** The direction of change in land use and coverage, according to classes and scenarios between 2000 and 2050. ↗ = *Increaseand* ↙.= *Reduction*.

|  | Forest veg-<br>etation | Grassland<br>vegetation | Planted<br>pasture | Agriculture | Mosaic of<br>occupation | Forestry |
| --- | --- | --- | --- | --- | --- | --- |
| Sustainable development<br>scenario<br>SSP1 RCP 1.9 | ↗ | ↘ | ↘ | ↗ | ↗ | ↘ |
| Middle of the road<br>scenario<br>SSP2 RCP 4.5 | ↘ | ↘ | ↗ | ↗ | ↗ | ↗ |
| Strong inequality<br>scenario<br>SSP3 RCP 7.0 | ↘ | ↘ | ↗ | ↗ | ↗ | ↗ |

According to the Middle of the road and Strong inequality scenarios, Forest vegetation will suffer a reduction of approximately 805,956 and 933,092 km^2^, respectively, until 2050, mainly in the Amazon biome (673,066 and 762,739 km^2^), followed by the Cerrado biome (67,643 and 87,288 km^2^). However, in the Sustainable development scenario, Forest vegetation will increase, occupying 447,944 km^2^. This increase will occur, mostly, in the Atlantic forest (253,783 km^2^), Cerrado (200,489 km^2^), and Caatinga (168,572 km^2^) biomes, as seen in Fig 4. While, only the Amazon biome will be reduced by 217,696 km^2^ of Forest vegetation, approximately 2/3 less than the values observed in the other scenarios. As seen in Fig 4 and Table 5, there will be a reduction in Grassland vegetation areas in both scenarios. Overall, in the Strong inequality scenario, the reduction will be greater with 1,346,988 km^2^ approximately, followed by the Middle of the Road scenario with 1,107,235 km^2^ and the Sustainable development scenario with 602,847 km^2^. Equivalent to what will occur with Forest vegetation, this reduction will occur mainly in the Cerrado (586,575 km^2^, on average) and Caatinga (303,419 km^2^, on average) biomes. Planted pasture and Forestry, inverse to what will happen with Forest vegetation, will tend to increase their extension in the Middle of the road and Strong inequality scenarios (219,199 and 439,962 km^2^, respectively), while in the Sustainable development scenario, there will be a reduction of, approximately 308.206 km^2^. Despite the data showing an increase in Agriculture and Mosaic of occupation, regardless of the scenario considered, the increase will occur with greater intensity in the Middle of the road and Strong inequality scenarios, whereas in Agriculture the increase will be 259,892 and 303,781 km^2^, respectively, and in the Mosaic of occupation this increase will be 1,366,687 and 1,450,867 km^2^, respectively. Although smaller, the increase in the Sustainable development scenario will correspond to 34,973 km^2^ in Agriculture and 403,914 km^2^ in Mosaic of occupation. It should be noted that the increase in the areas of Planted pasture, Forestry, and Agriculture, should occur in the Cerrado biome, while the increase in the Mosaic of occupation will occur, mostly in the Amazon and Cerrado biomes. This set of scenarios provides important information that can corroborate efforts to establish public policies that aim to contribute to the conservation of biodiversity and reduce emissions from deforestation and degradation, especially those arising from changes in land use and coverage. Furthermore, this set of scenarios with territorial extension for the whole of Brazil, makes it possible to understand how decision-making and global demands, in some determined regions, can influence other regions.

## Supporting information

**S1 Appendix. LuccMEBR: scenario-dependent spatiotemporal drivers**.

**S2 Appendix. LuccMEBR: Model parameters**.

## Acknowledgments

The authors thank the project “MSA / BNDES (Environmental Monitoring by Satellite in the Amazon biome)” for financing the development of LuccMEBR. We also thank Eloi Dalla Nora and Detlef Van Vuuren for their contribution to the development of the scenarios.

